# *TCF7L2* is a master regulator of insulin production and processing

**DOI:** 10.1101/003202

**Authors:** Yuedan Zhou, Soo Young Park, Jing Su, Kathleen Bailey, Emilia Ottosson-Laakso, Liliya Shcherbina, Nikolay Oskolkov, Enming Zhang, Thomas Thevenin, João Fadista, Hedvig Bennet, Petter Vikman, Nils Wierup, Malin Fex, Johan Rung, Claes Wollheim, Marcelo Nobrega, Erik Renström, Leif Groop, Ola Hansson

## Abstract

Although variants in the T-cell factor 7-like 2 gene (*TCF7L2*) confer the strongest risk of type 2 diabetes (T2D) by presumed effects on islet function, the underlying mechanisms are not well understood. We have identified TCF7L2-target genes and described the regulatory network downstream of TCF7L2 responsible for its effect on insulin secretion in rodents and human pancreatic islets. *ISL1* is a direct target of TCF7L2 and regulates proinsulin production and processing via *MAFA*, *PDX1*, *NKX6.1*, *PCSK1* and *PCSK2* and possibly clearance of proinsulin via *SLC30A8*. Taken together, these results demonstrate that not only synthesis of proinsulin is regulated by TCF7L2, but also processing and possibly clearance of proinsulin and insulin in a genotype dependent manner. These multiple targets in key pathways may explain why *TCF7L2* has emerged as the gene showing the strongest association with T2D.

## Introduction

*TCF7L2* harbours genetic variants with the strongest effect on T2D risk yet described, *i.e.* the single nucleotide polymorphism (SNP rs7903146)(Saxena et al., 2012). TCF7L2 is a transcription factor (TF) in the Wnt-signalling pathway and expressed in many tissues including fat, liver and pancreatic islets of Langerhans(Osmark et al., 2009). A majority of variants associated with T2D by GWAS seem to affect islet function(Florez, 2008). In concordance, the risk T-allele of rs7903146 is associated with impaired glucose-stimulated insulin secretion (GSIS) or other secretagogues like GLP-1(Chang et al., 2010; Loder et al., 2008; Lyssenko et al., 2007; Rosengren et al., 2012; Saxena et al., 2006; Shu et al., 2009; Shu et al., 2008) Risk T-allele carriers are further characterized by an elevated plasma proinsulin level and an increased proinsulin/insulin ratio suggestive of perturbed proinsulin processing(Dahlgren et al., 2007; Dupuis et al., 2010; Gonzalez-Sanchez et al., 2008; Kirchhoff et al., 2008; Loos et al., 2007; McCaffery et al., 2011; Silbernagel et al., 2011; Stolerman et al., 2009; Strawbridge et al., 2011). In support of a primary effect in pancreatic islets, the risk T-allele carriers show higher degree of open chromatin in pancreatic islets, but not in other tissues(Gaulton et al., 2010). Several mouse models of *Tcf7l2* have been investigated(Boj et al., 2012; da Silva Xavier et al., 2012; Korinek et al., 1998; Savic et al., 2011b; Takamoto et al., 2013; Yang et al., 2012), but the diabetic phenotype has not been replicated in all studies. Experiments in rodent islets where TCF7L2 activity has been disrupted have usually demonstrated impaired GSIS(da Silva Xavier et al., 2009; Shu et al., 2008; Zhou et al., 2012).

Paradoxically, carriers of the risk T-allele show increased expression of *TCF7L2* in human pancreatic islets(Le Bacquer et al., 2012; Lyssenko et al., 2007). The T-allele locus displays much stronger transcriptional activity(Gaulton et al., 2010; Stitzel et al., 2010). TCF7L2 may act both as a stimulator and a repressor of gene expression due to both differential splicing and interaction with different co-regulators in target gene recognition(Hansson et al., 2010). However, the target genes of TCF7L2 in pancreatic islets or β-cells have not been described in detail. Our aim was to identify TCF7L2 target genes in pancreatic islets and to describe the regulatory network downstream of TCF7L2 responsible for its influence on insulin secretion.

## Results

### Tcf7l2 influences insulin synthesis and secretion

Impaired *in vivo*(Lyssenko et al., 2007) and *in vitro* (Le Bacquer et al., 2012; Rosengren et al., 2012) insulin secretion has been shown in risk T-allele carriers of rs7903146. We have replicated and extended these findings in pancreatic islets from human non-diabetic cadaver donors (n=75, Figure 1A and Supplemental Table S3). We also show reduced insulin content (27%) in risk T-allele carriers (n=81, Figure 1B), and that *TCF7L2* mRNA expression is 16% higher than C-allele carriers (n=66; RNA-seq, Figure 1C). Further, *TCF7L2* expression is negatively associated with proinsulin expression in human islets in C-allele carriers (n=36), but not in T-allele carriers (n=30, Figure 1D, Table S4).

**Figure 1.**
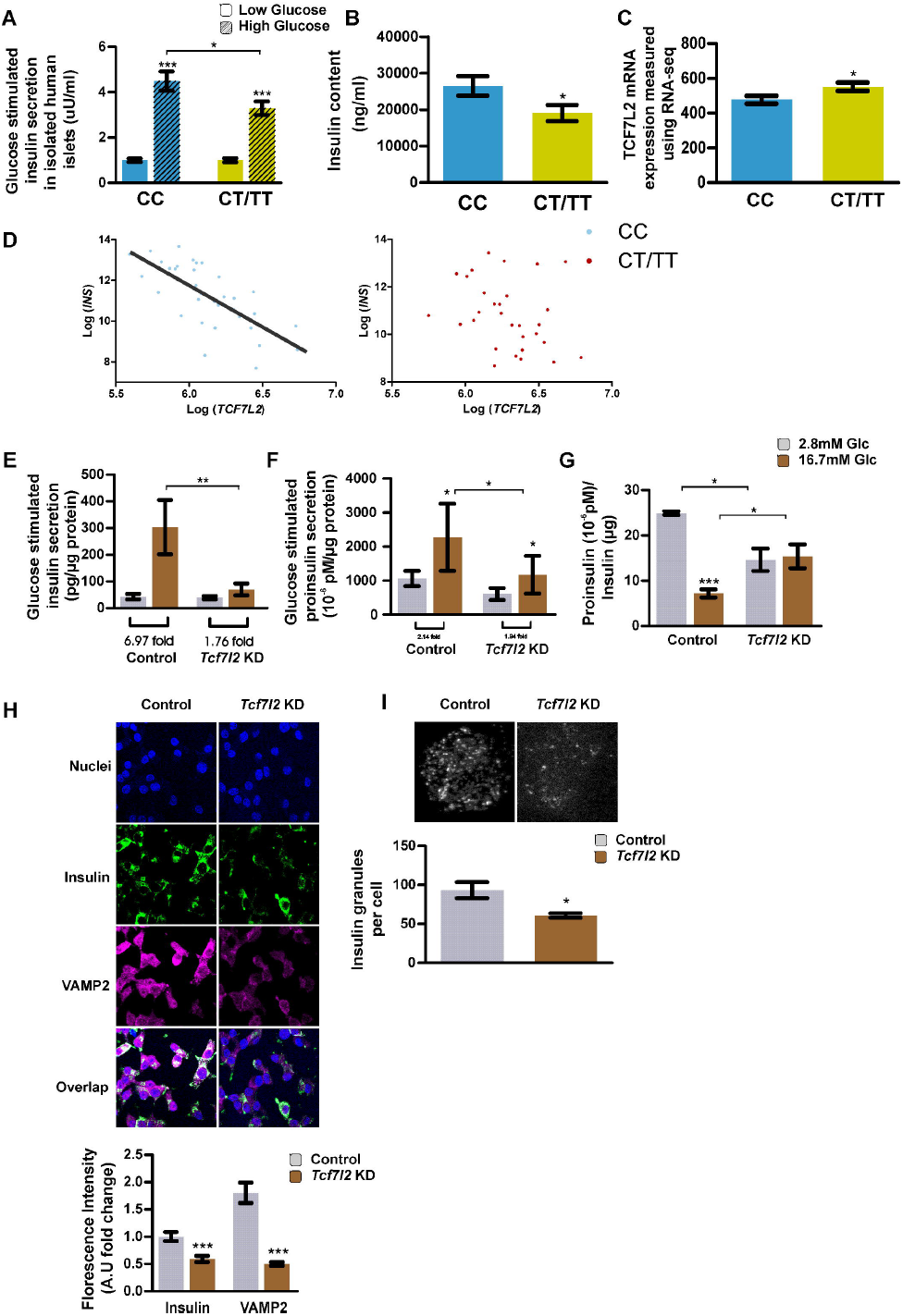
The type 2 diabetes associated SNP rs7903146 and *TCF7L2* expression influences insulin synthesis and secretion. (A) Glucose-stimulated insulin secretion in human islets (CC: n = 44, CT/TT: n = 31) (B) Insulin content in human islets (CC: n = 45, CT/TT: n = 36). (C) *TCF7L2* mRNA expression measured by RNA-seq in human islets (CC: n = 36, CT/TT: n = 30) (D) Expression (mRNA) of *INS* and *TCF7L2* in human islets measured by RNA-seq stratified for rs7903146 genotype (CC: n = 36, CT/TT: n = 30). (E) Glucose-stimulated insulin secretion in INS1 cells after *Tcf7l2* silencing. (F) Glucose-stimulated proinsulin secretion in INS1 cells after *Tcf7l2* silencing. (G) Proinsulin-to-insulin ratio in INS1 cells after *Tcf7l2* silencing. (H) Immunohistochemistry staining of insulin and Vamp2 in INS1 cells after *Tcf7l2* silencing. (I) Amount of insulin measured by TIRF microscopy in INS1 cells after *Tcf7l2* silencing. Data are represented as mean ± SEM. In (A-D) data were analyzed by linear regression with correction for age, gender and BMI. In (E-I), 2.8 versus 16.7 mM glucose were analyzed by paired Student’s t-tests, Tcf7l2 silencing versus Controls by unpaired Student’s t-tests with n = 3. * = p < 0.05, ** = p < 0.01, *** = p < 0.001, (Table S1 and S4).

Silencing *Tcf7l2* in a rat insulinoma cell line (INS1 832/13) using siRNA targeting the invariable exon 1, resulted in 84% knock-down of *Tcf7l2* mRNA, 90% reduction of TCF7L2 activity and a reduced expression of the well-known TCF7L2 target genes *Axin2* and *Ccnd1* (Figure S1A-B)(Zhou et al., 2012). The decrease in *Tcf7l2* resulted in a marked reduction of GSIS, *i.e.* 1.76 fold (*Tcf7l2* silencing, p>0.05) versus 6.97 fold (controls, p<0.05, Figure 1E) and a perturbed glucose-stimulated proinsulin secretion (GSPS) (p<0.05, Figure 1F). Proinsulin/insulin ratio was higher under glucose stimulation compared to basal condition in *Tcf7l2* silenced cells (p<0.05, Figure 1G) suggesting perturbed proinsulin processing. Immunohistochemistry (IHC) analysis of *Tcf7l2* silenced INS1 cells showed reduced staining intensity for insulin (p<0.00001) and VAMP2 (p<0.00000001), a marker of insulin-containing granules, in *Tcf7l2* silenced cells (Figure 1H). Furthermore, there was a reduction in insulin containing granules close to the plasma membrane as assessed by total internal reflection fluorescence (TIRF) microscopy (p<0.05, Figure 1I). Together, these data suggest that TCF7L2 mediates key aspects of insulin production, processing and intracellular trafficking.

### Identification of Tcf7l2 target genes influencing insulin synthesis and secretion

The molecular mechanisms underlying these findings are not clear, but an altered expression of TCF7L2-target genes is likely part of the explanation. To identify genes regulated by TCF7L2 in β-cells, we screened for global differential expression using RNA-seq in *Tcf7l2* silenced INS1 cells versus scramble-treated control. Overall, 10779 genes were actively expressed (>30 count per gene); 1680 genes were up-regulated and 1885 genes were down-regulated in *Tcf7l2* silenced cells (5% FDR, n=4). Among the most differentially expressed genes were *Il6r*, *Aldh1a1* and *Srp14*, but none of these has a known function in insulin secretion. Genetic variation in the *IL6R*-locus has been weakly associated with T2D (Hamid et al., 2004), but this was not replicated (Qi et al., 2007) or supported by recent GWAS findings (Cho et al., 2012; Dupuis et al., 2010; Kooner et al., 2011; Saxena et al., 2007; Scott et al., 2007; Sladek et al., 2007; Voight et al., 2010; Zeggini et al., 2007) therefore *Il6r* was not studied further. Instead we categorized genes known to influence GSIS into three groups: (i) glucose sensing, (ii) proinsulin expression and maturation and (iii) exocytotic machinery (Table 1). In addition, genes located proximal to SNPs with replicated associations to T2D and/or plasma proinsulin were also selected for further analysis (Table S2). Real-time quantitative PCR (QPCR) confirmed the RNA-seq results for many of the tested genes (Table 1), *i.e.* 18 out of 29 analyzed genes were replicated in independent samples at the mRNA level. To confirm these findings at the protein level, a selected set of 15 genes (with mRNA change >30%) was analyzed using Western blot, with 11 genes having significant changes (Table S1).

**Table 1.**
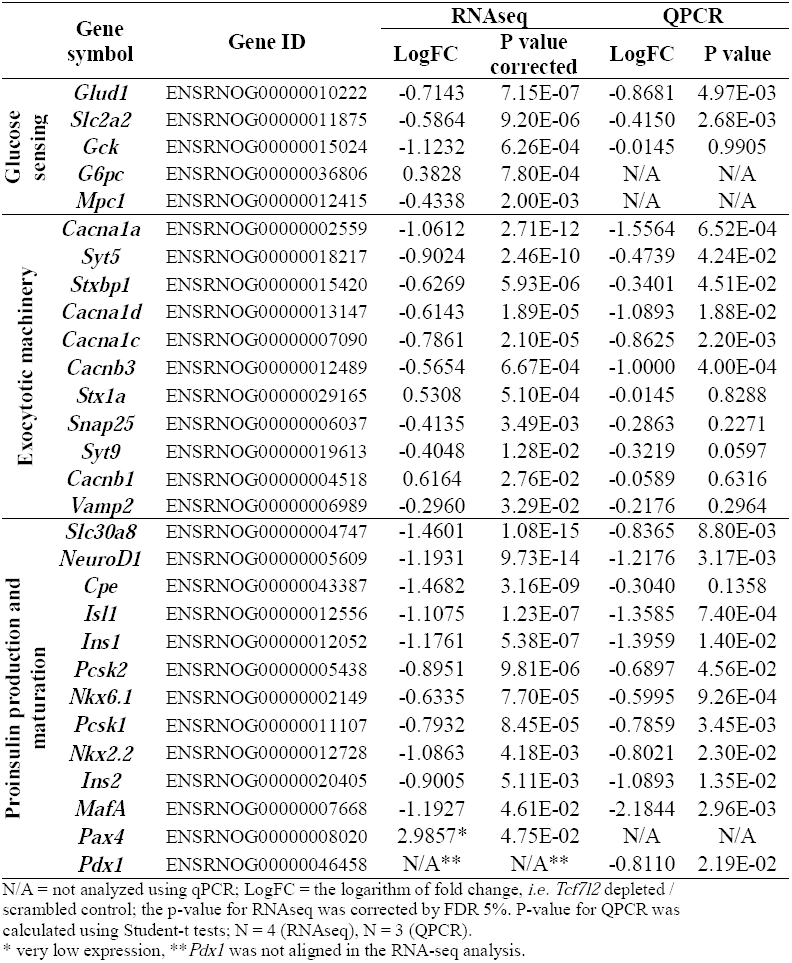
mRNA expression of selected genes influencing glucose stimulated insulin secretion measured using RNA sequencing (RNAseq) in *Tcf7l2* depleted INS1 832/13 cells versus scrambled control, with replication in independent experiments using real-time quantitative PCR (QPCR).

### Tcf7l2 is a key regulator of proinsulin production and processing

Twelve analyzed genes involved in proinsulin synthesis were down-regulated in *Tcf7l2* silenced cells, including proinsulin genes *Ins1* and *Ins2* (Table 1 and Figure 2A). The reduction of all but one (*Cpe*) was confirmed using QPCR. The protein encoded by the *Ins1* and *Ins2* genes was decreased in *Tcf7l2* silenced cells (Figure 2B). We next replicated this finding in primary cells by measuring proinsulin protein expression in a *Tcf7l2* homozygous null mouse model (*Tcf7l2*^-/-^)(Savic et al., 2011b). Notably proinsulin expression was reduced by 85% (n=6) in pancreas from *Tcf7l2*^-/-^ mice (postnatal day 0; P0) compared to wild type mice (Figure 2C). Given differences in the insulin gene between rodents and humans(Soares et al., 1985), we also replicated the findings in human pancreatic islets showing 49% reduced proinsulin mRNA expression after *TCF7L2* silencing (n=5, Figure 2D).

**Figure 2.**
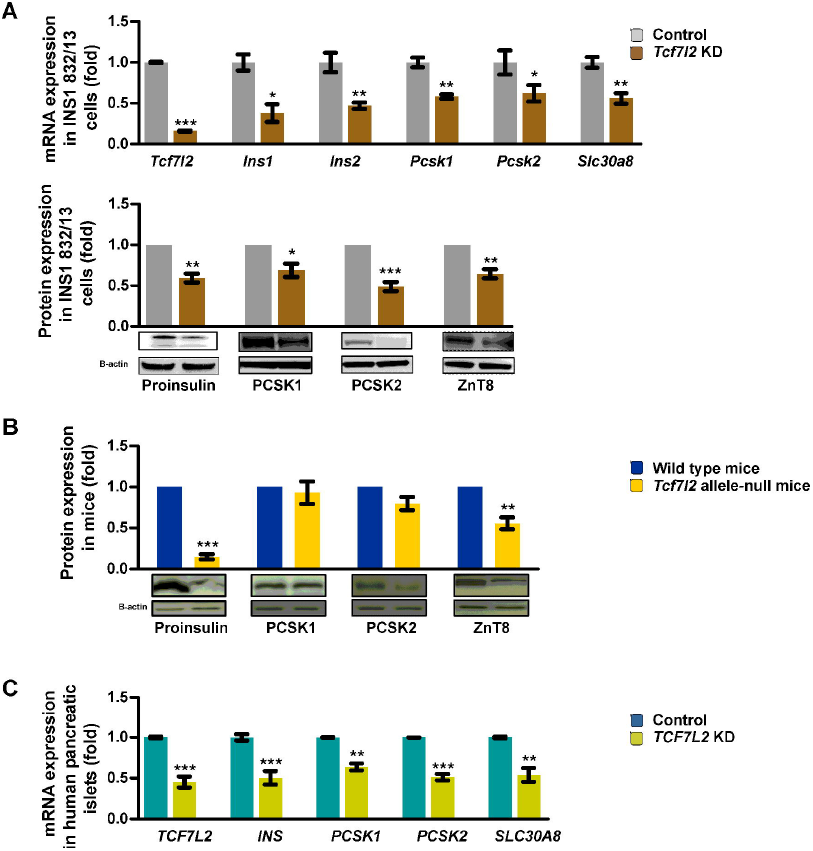
TCF7L2 is a key-regulator of proinsulin production and processing. (A) mRNA expression measured by QPCR in INS1 cells in *Tcf7l2* silenced- versus control cells, (n = 3). Protein expression measured by Western blot in INS1 cells in *Tcf7l2* silenced- versus control cells, (n = 3). (B) Protein expression measured by Western blot in *Tcf7l2* allele-null versus wild type mice (n = 6). (C) mRNA expression measured by QPCR in human pancreatic islets in *Tcf7l2* silenced- versus control cells, (n = 3 – 5). Data are represented as mean ± SEM. * = p < 0.05, ** = p < 0.01, *** = p < 0.001 analyzed by Student’s t-tests, (Table S1).

In the processing of proinsulin-to-insulin, the endoprotease prohormone convertase 1/3 (PCSK1) and prohormone convertase 2 (PCSK2) cleave the proinsulin molecule to yield mature insulin and C-peptide. We found reduced mRNA expression levels of both *Pcsk1* and *Pcsk2* in the RNA-seq screen of *Tcf7l2* silenced INS1 cells (Table 1). This result was confirmed using QPCR and at the protein level in INS1 cells (Figure 2A-B). No significant change in PCSK1 and PCSK2 protein expression was observed in the *Tcf7l2*^-/-^ mouse pancreas (Figure 2C). In *TCF7L2* silenced human pancreatic islets, *PCSK1* and *PCSK2* mRNA expression was reduced by 36% and 49% respectively (Figure 2D).

### Tcf7l2 is not a major regulator of genes involved in the exocytotic machinery or glucose sensing

Of the 11 genes categorized as involved in the exocytotic machinery, mRNA expression of *Cacna1a*, *Cana1c*, *Cacna1d* and *Cacnb3* was reduced in *Tcf7l2* silenced cells (Table 1 and Supplemental Figure S1C). However, this was not confirmed at the protein level for Cav2.1, Cav1.2 or Cav1.3, either 72 hours, or 7 days after *Tcf7l2* silencing (Figure S1D-E). RNA expression of 2 out of the 5 genes involved in glucose sensing (*Glud1* and *Slc2a2/Glut2*) was reduced at the mRNA level in *Tcf7l2* silenced cells (Table 1), and GLUD1 protein expression was measured but no change was observed (Figure S1F).

### Tcf7l2 is regulating proinsulin expression directly via Isl1, Ins1 and Ins2 and indirectly via MafA, Neurod1 and Pdx1

After establishing that TCF7L2 is a regulator of proinsulin expression and maturation, we investigated by which molecular mechanism this may occur. For this purpose, we interrogated the RNA-seq results for genes differentially expressed between *Tcf7l2* silenced INS1 and control cells. Intriguingly, ISL LIM homeobox 1 (*Isl1*), v-maf avian musculoaponeurotic fibrosarcoma oncogene homolog A (*MafA*), pancreatic and duodenal homeobox 1 (*Pdx1*), neuronal differentiation 1 (*NeuroD1*), NK6 homeobox 1 (*Nkx6.1*) and NK2 homeobox 2 (*Nkx2.2*) were all expressed at lower levels in *Tcf7l2* silenced cells (Table 1). All observations, except the NKX2.2 protein, were confirmed using QPCR and Western blot (Figure 3A). The reduction of ISL1 and NEUROD1 was further confirmed using IHC (Figure S1G-H). These results were confirmed in *Tcf7l2*^-/-^ mice, with protein levels reduced relative to wild type mice for ISL1, MAFA and NEUROD1 (Figure 3B). Likewise, after *TCF7L2* silencing in human islets, expression of most transcripts was down-regulated, including *ISL1*, *MAFA*, *PDX1*, *NEUROD1* and *NKX6.1* (Figure 3C). In addition, in islets from CC-genotype carriers of rs7903146, *TCF7L2* mRNA expression was negatively associated with that of *ISL1*, *MAFA* and *NKX6.1* but not for *MAFA* and *NKX6.1* in CT/TT-risk-genotype carriers (Table S4). The disruption of *Tcf7l2* resulted in a marked reduction in GSIS compared with controls 1.93 (p>0.05) *vs.* 4.10 fold (p<0.05) increases (Figure 3D). The same was observed after disruption of *Isl1*, *MafA* and *NeuroD1* - (Figure 3D).

**Figure 3.**
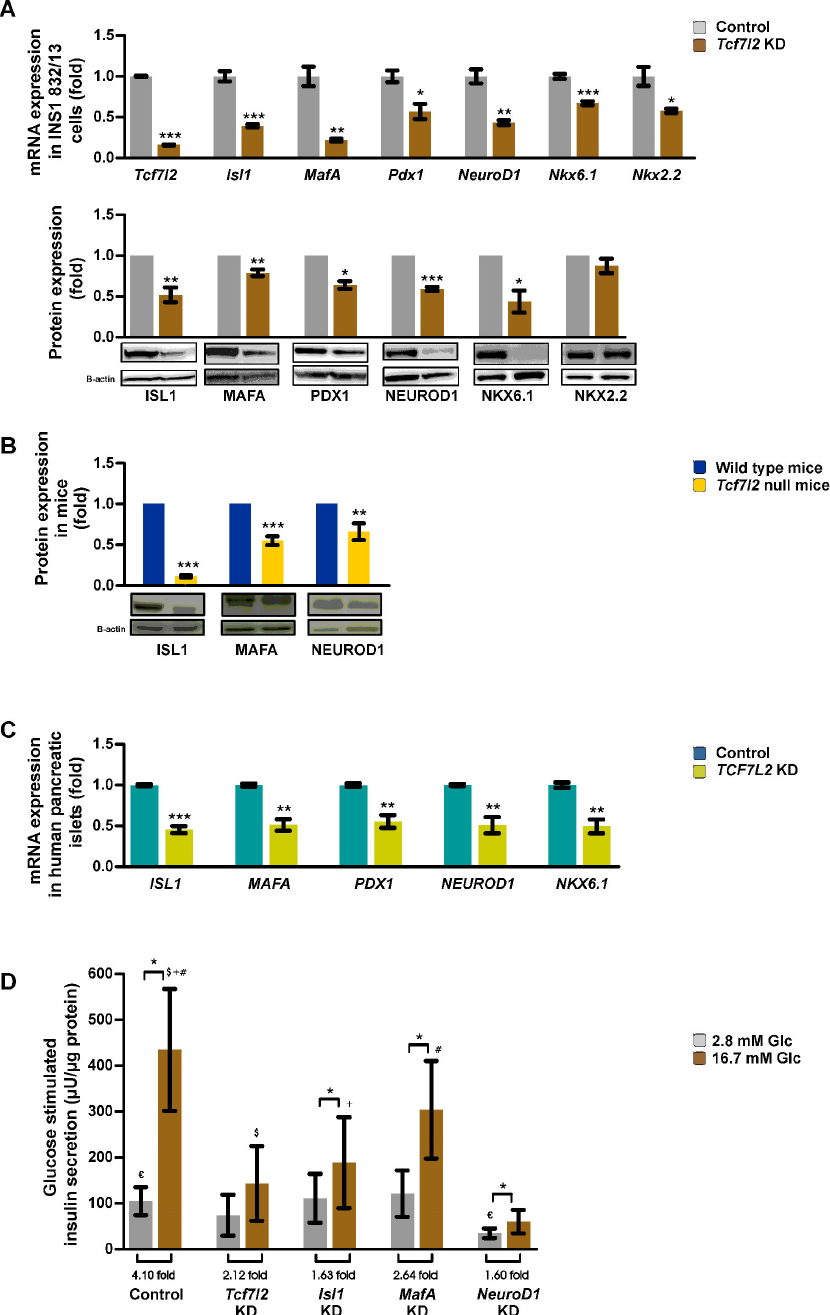
TCF7L2 is regulating proinsulin expression via direct regulation of ISL1 and indirectly via MAFA, NEUROD1 and PDX1. (A) mRNA expression measured by QPCR in INS1 cells in *Tcf7l2* silenced- versus control cells, (n = 3). Protein expression measured by Western blot in INS1 cells in *Tcf7l2* silenced- versus control cells, (n = 3). (B) Protein expression measured by Western blot in *Tcf7l2* allele-null versus wild type mice (n = 6). (C) mRNA expression measured by QPCR in human pancreatic islets in *Tcf7l2* silenced- versus control cells, (n = 3). (D) Glucose-stimulated insulin secretion in INS1 cells after *Tcf7l2*, *Isl1*, *MafA* and *NeuroD1* silencing, (n = 3). Data are represented as mean ± SEM. * = p < 0.05, ** = p < 0.01, *** = p < 0.001 analyzed in (A – C) by unpaired Student’s t-tests. In (D), 2.8 versus 16.7 mM glucose were analyzed by paired Student’s t-tests (Table S1).

To delineate the proinsulin regulatory cascade among the identified TCF7L2 target genes, *Isl1*, *MafA*, *Pdx1*, *NeuroD1* and *Nkx6.1* were sequentially silenced in INS1 cells using siRNA. First, knocking down *Isl1* reduced both the mRNA and protein expression of *MafA*, *Pdx1*, *Nkx6.1*, *Pcsk1*, *Pcsk2*, *Ins1* and *Ins2* (Figure 4A). No effect of *Isl1* knock-down was observed on *Tcf7l2* or *NeuroD1* mRNA expression (Figure 4A). Previously we have analysed chromatin immunoprecipitation on tiling array (ChIP-on-Chip) using a TCF7L2 antibody identifying both the *Isl1* and *Ins1* promoters as directly bound by TCF7L2 (Zhou et al., 2012). Together, these data indicate that *Isl1* is a direct primary target gene for TCF7L2 and regulates proinsulin expression and processing via *Isl1* dependent regulation of *MafA*, *Pdx1*, *Nkx6.1*, *Pcsk1* and *Pcsk2*. In human islets, *ISL1* expression is positively associated in C-allele carriers with that of *MAFA*, *PDX1*, *NKX6.1*, *INS* and *SLC30A8* but in T-allele carriers only with *PDX1* and *NKX6.1* (Table S4).

**Figure 4.**
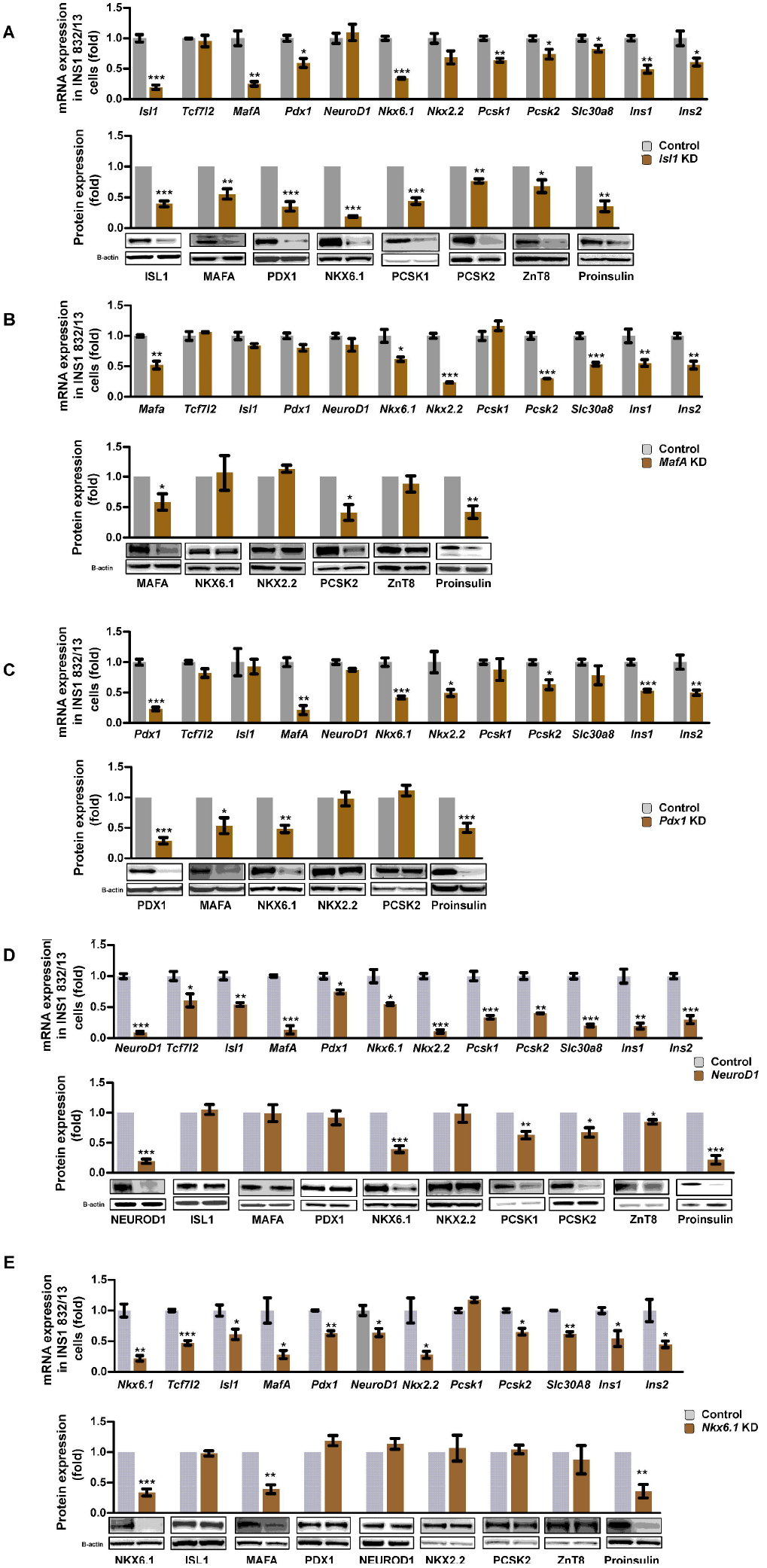
Sequential silencing of TCF7L2 target genes in the proinsulin regulatory cascade. mRNA and protein expression after: *Isl1* (A), *MafA* (B), *Pdx1* (C), *NeuroD1* (D) and *Nkx6.1* (E) silencing versus controls measured by QPCR and Western blot in INS1 cells. Data are represented as mean ± SEM. * = p < 0.05, ** = p < 0.01, *** = p < 0.001 analyzed by Student’s t-tests with n = 3, (Table S1).

Knocking down *MafA* reduced mRNA expression of *Nkx6.1*, *Nkx2.2*, *Pcsk2*, *Ins1* and *Ins2*. These changes were confirmed at the protein level for all but NKX6.1 and NKX2.2 (Figure 4B). In human islets, *MAFA* expression was positively associated with proinsulin gene expression both in C- and T-allele carriers (Table S4). Knocking down *Pdx1* reduced the mRNA expression of *MafA*, *Nkx6.1*, *Nkx2.2*, *Pcsk2*, *Ins1* and *Ins2*. These changes were confirmed at the protein level for all but PCSK2 (Figure 4C). Knocking down *NeuroD1* reduced the mRNA expression of *Tcf7l2*, *Isl1, MafA*, *Pdx1, Nkx6.1*, *Nkx2.2, Pcsk1*, *Pcsk2*, *Ins1* and *Ins2*. This was, however, not reflected by changes in the ISL1, MAFA and PDX1 protein levels (Figure 4D). Knocking down *Nkx6.1* reduced the mRNA expression of *Tcf7l2*, *Isl1*, *MafA*, *Pdx1*, *Nkx6.1*, *Nkx2.2*, *Pcsk2*, *Ins1* and *Ins2*. At the protein level, we could only confirm the reduction of MAFA and proinsulin (Figure 4E).

In summary, these data together with previous ChIP-on-Chip results (Zhou et al., 2012) indicate that ISL1 is a direct target of TCF7L2 and that ISL1, in turn regulates proinsulin production and processing via regulation of MAFA, PDX1, NKX6.1, PCSK1 and PCSK2 expression.

### TCF7L2 is a transcriptional regulator of key genes located in T2D-associated loci

Next we examined whether silencing of *Tcf7l2* was associated with changes in expression of rat orthologues of putative genes in T2D–associated loci (Cho et al., 2012; de Miguel-Yanes et al., 2011; Kooner et al., 2011; Morris et al., 2012; Saxena et al., 2012; Voight et al., 2010)(Table S2). We observed differential mRNA expression for 23 genes, 17 out of these were replicated using QPCR and 3 at the protein level (*i.e. Slc30a8*, *Ide* and *Bcl11a*)-(Figure 2 and 5). Of note, *Tcf7l2* silencing in INS1 cells led to a decreased mRNA expression of *Slc30a8* (Figure 2A). In pancreata from *Tcf7l2*^-/-^ mice, ZnT8 (*Slc30a8*) expression was reduced by 44% compared to wild type control mice (n=6, Figure 2C). This was also replicated in human islets showing a 46% reduction of *SLC30A8* mRNA expression after *TCF7L2* silencing (Figure 2D). ZnT8 protein expression was also reduced in INS1 cells after *Isl1* and *NeuroD1* silencing (Figure 4A-D), but not after *Pdx1, MafA* and *Nkx6.1* silencing (Figure 4B, C and E).

**Figure 5.**
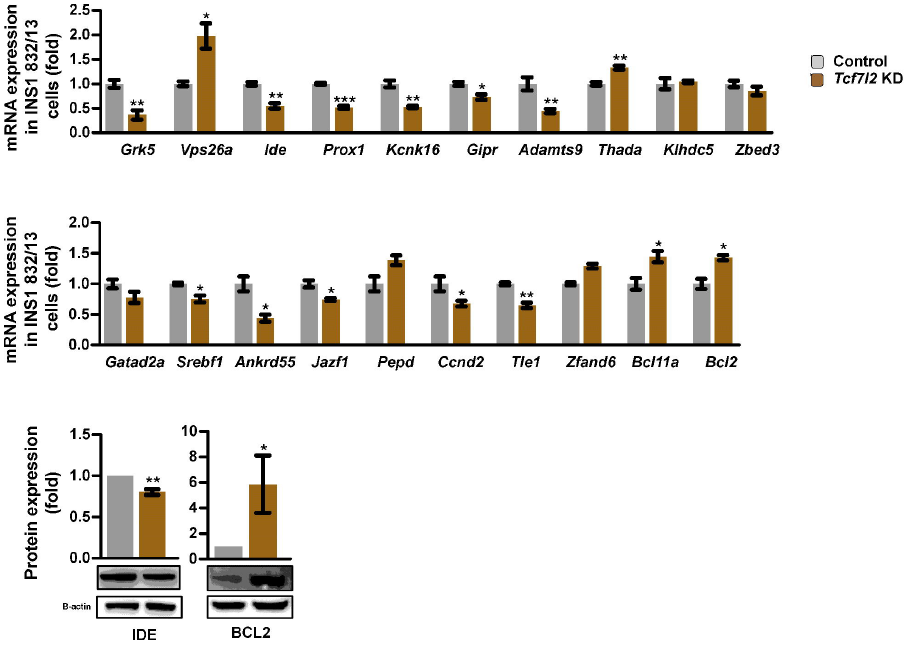
TCF7L2 is a transcriptional regulator of key genes located in T2D-associated loci. mRNA expression measured by QPCR in INS1 cells in *Tcf7l2* silenced- versus control cells, (n = 3). Protein expression was measured for two genes, *i.e.* IDE and BCL2, by Western blot in INS1 cells in *Tcf7l2* silenced- versus controls, (n = 3). Data are represented as mean ± SEM. * = p < 0.05, ** = p < 0.01, *** = p < 0.001 analyzed by Student’s t-tests, (Table S1).

Finally, expression (mRNA) of *Grk5*, *Ide*, *Prox1*, *Kcnk16*, *Gipr*, *Adamts9*, *Srebf1*, *Ankrd55*, *Jazf1*, *Ccnd2* and *Tle1* was down-regulated whereas expression of *Vps26a*, *Thada*, *Bcl11a* and *Bcl2* was up-regulated after *Tcf7l2* knock-down (Table S2). In addition, the BCL11A protein expression was markedly up-regulated after disruption of *Isl1* and *NeuroD1* (Figure 4A and D).

## Discussion

The current report presents a genetic network controlled by TCF7L2 in pancreatic islets, providing a comprehensive view of the molecular mechanisms by which TCF7L2 regulates glucose metabolism. We demonstrate that TCF7L2 plays a central role in coordinating the expression of proinsulin and its subsequent processing to form mature insulin. We also show that TCF7L2 and ISL1 are key regulators of this transcriptional network and provide evidence that they regulate MAFA, PDX1, NEUROD1 and NKX6.1 (Figure 6). The two prohormone convertases responsible for the processing of proinsulin to mature insulin, were also targets of TCF7L2/ISL1-regulation. These functions seem to have been highly conserved during evolution as similar results were obtained in a rat cell line, primary mouse and human islets. TCF7L2 is also regulates several genes in T2D-associated loci. Of note, *SLC30A8* (encoding the zinc transporter ZnT8) is a downstream target of TCF7L2/ISL1-regulation. Zinc is important for the formation of insulin crystals (Scott, 1934), and a common non-synonymous SNP in the zinc transporter *SLC30A8* has been associated with both T2D (Sladek et al., 2007) and reduced proinsulin-to-insulin conversion (Kirchhoff et al., 2008). *SLC30A8* has previously been identified as a TCF7L2-target gene in mice (da Silva Xavier et al., 2009), a finding now replicated in humans. Although the mechanisms by which the variant in *SLC30A8* increases risk of T2D are not clear, recent data suggest that reduction in zinc concentrations in the portal vein after β-cell specific disruption of *Slc30a8* might influence hepatic insulin clearance (Tamaki et al., 2013). If confirmed, this would suggest that *TCF7L2* might influence all key steps in (pro)insulin synthesis, processing and clearance.

**Figure 6.**
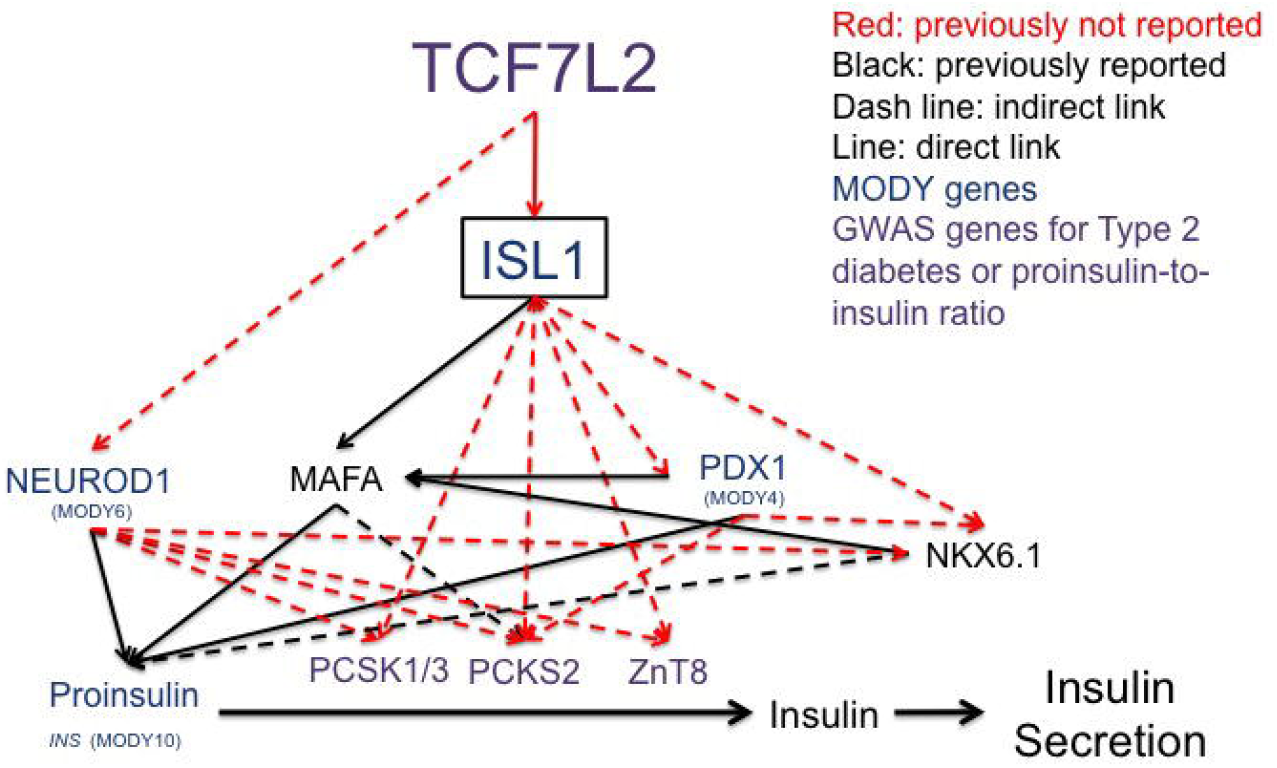
Schematic figure of how TCF7L2 functions as a master regulator of insulin production and processing. Schematic figure showing identified TCF7L2-target genes and the downstream regulatory network responsible for its effect on insulin secretion. In rodents, as well as human pancreatic islets, *ISL1* is a direct target of TCF7L2. ISL1 regulates proinsulin production and processing via regulation of PCSK1, PCSK2, SLC30A8, MAFA, PDX1 and NKX6.1. These multiple targets in key pathways may explain why *TCF7L2* has emerged as the gene showing the strongest association with T2D.

Transcription of proinsulin in humans is under the control of many glucose responsive TFs that mainly bind a 400 base pair conserved region (Fu et al., 2013; Hay and Docherty, 2006). Among the most studied TFs are MAFA, PDX1 and NEUROD1, all shown here to be transcriptional targets of TCF7L2/ISL1 regulation. In support of this, ISL1 regulates *MAFA* expression by directly binding to the *MAFA* promoter (Du et al., 2009), but its influence on *Pdx1*, *Nkx6.1*, *Pcsk1* and *Pcsk2* expression has not been described to our knowledge. Together with our recent demonstration of direct binding of TCF7L2 to the *Isl1* promoter (Zhou et al., 2012), these data indicate that TCF7L2 together with ISL1 forms a master regulatory circuit influencing both proinsulin expression and processing via regulation of *MafA*, *Pdx1*, *Nkx6.1*, *Pcsk1* and *Pcsk2*. As ISL1 also can bind and influence the expression of other pancreatic hormones like glucagon (Wang and Drucker, 1995) and somatostatin (Leonard et al., 1992), this indicates that TCF7L2 may also have important regulatory functions in other islet cell types besides the β-cell. Relatively high expression of *TCF7L2* in α-cells has also been reported in sorted human pancreatic islet cells (Kirkpatrick et al., 2010). A recent study indicated that NEUROD1 regulates the expression of *Ins1* but not *Ins2* in primary mouse β-cells (Gu et al., 2010), while we observed altered expression of both proinsulin genes as well as *Nkx6.1* in rat cell line.

TCF7L2 may also regulate other β-cell functions essential for maintaining GSIS. A perturbation of intracellular [Ca^2+^] was observed in one *Tcf7l2*^-/-^ mouse model, but no major effect on the expression of Ca^2+^ channels (da Silva Xavier et al., 2009). Here, we find that although the mRNA levels of several Ca^2+^ channels important for GSIS were reduced in *Tcf7l2* silenced cells, no changes of the corresponding proteins were observed either 72 hours or 7 days after *Tcf7l2* knock-down. An effect of TCF7L2 on Ca^2+^ handling in the β-cell should however not be excluded based upon these data as these proteins may have a very long half-life in the cell. An influence on cell viability and regulation of apoptosis are other well-documented effects of TCF7L2 in pancreatic islets (Le Bacquer et al., 2011; Shu et al., 2008; Zhou et al., 2012). The T2D-associated SNP rs7903146 has also been associated with pancreatic islet morphology, *i.e*. with reduced islets density, reduction of islet size and an increased α- to β-cell ratio in risk-genotype carriers (Le Bacquer et al., 2012). In the same report, an association between the rs7903146 and GSIS was seen; but not with insulin content per islet equivalent as shown in our study. There may be several explanations for these discrepancies but a large number of high quality human islets are probably needed to demonstrate a significant reduction in insulin content, as the state of the islets can have a large influence on insulin measurements.

Some recent reports have questioned the dogma that genetic variants in *TCF7L2* exert its major effect on islets (Boj et al., 2012; Shao et al., 2013). Boj *et al* recently reported reduced hepatic glucose production in a liver specific *Tcf7l2* knock out mouse model (Boj et al., 2012). We and others have shown an association between the rs7903146 variant and impaired suppression of hepatic glucose production (Lyssenko et al., 2007; Pilgaard et al., 2009; Wegner et al., 2008), but it is not easy to disentangle these effects from the effects of a reduction in insulin secretion (Faerch et al., 2013). Surprisingly, they reported no impaired effects on β-cell function in a β-cell specific *Tcf7l2* knockout mouse (Boj et al., 2012). This finding is in sharp contrast with previous reports showing reduced β-cell mass, a perturbed incretin response and GSIS with reduced insulin expression (da Silva Xavier et al., 2012; Takamoto et al., 2013). Although there are no apparent explanations for these discrepant results they could reflect the application of β-cell specific versus whole-islet knock-down strategies, as *TCF7L2* is also expressed in α-cells (Kirkpatrick et al., 2010), or a difference between inducible versus congenital knock-down of *TCF7L2*, as Wnt-signaling is important for pancreas development (Murtaugh, 2008). An alternative explanation is a possible compensatory increased expression of other genes taking over the role of TCF7L2 in the β-cells, in support is the reported lack of regulation of well-known TCF7L2 target genes, *i.e.* Axin2 and Sp5 (Boj et al., 2012). TCF7L2 is expressed in many tissues besides the pancreas and there is no doubt that it has central functions in tissues like the brain (Lee et al., 2009; Savic et al., 2011a; Shao et al., 2013), liver (Boj et al., 2012; Ip et al., 2012; Oh et al., 2012), adipose tissue (Mondal et al., 2010; Prokunina-Olsson et al., 2009) and the gut (Kaminska et al., 2012; Shao et al., 2013). However, given the central role of insulin for maintenance of normal glucose tolerance it is unlikely that an extrapancreatic mechanism could exert an effect size of that seen for the rs7903146 in *TCF7L2*.

Interestingly, in human islets we observe a negative association between *TCF7L2* expression and target gene expression in the proinsulin synthesis pathway (Figure 1D), but also find that silencing total expression of *TCF7L2* leads to reduced proinsulin expression (Figure 2C). Experimental disruption of TCF7L2 activity has previously been associated with perturbed islet function (da Silva Xavier et al., 2009; da Silva Xavier et al., 2012; Le Bacquer et al., 2012; Shu et al., 2009; Shu et al., 2008; Zhou et al., 2012) but the risk T-allele is suggested to lead to increased expression of *TCF7L2* (Figure 1C)(Gaulton et al., 2010; Le Bacquer et al., 2012; Lyssenko et al., 2007; Stitzel et al., 2010) and perturbed GSIS in human islets (Figure 1A)(Chang et al., 2010; Le Bacquer et al., 2012; Loder et al., 2008; Lyssenko et al., 2007; Rosengren et al., 2012; Saxena et al., 2006; Shu et al., 2009; Shu et al., 2008). These seemingly contradictory findings might reflect that total expression of *TCF7L2* do not mirror TCF7L2 activity and rather it is the ratio of activating and inhibiting TCF7L2 isoforms that determine the activity and hence target gene regulation (Hansson et al., 2010).

In summary, in rodents as well as human pancreatic islets, *ISL1* is a direct target of TCF7L2 and ISL1, in turn, regulates proinsulin production and processing via regulation of PCSK1, PCSK2, SLC30A8, MAFA, PDX1 and NKX6.1. Furthermore, TCF7L2 might also influence hepatic clearance of insulin via its effect on SLC30A8. Taken together the current results provide some explanations for the large impact of *TCF7L2* on the pathogenesis of T2D demonstrating that TCF7L2 is a key regulator of (pro)insulin synthesis, processing and possibly clearance.

## Methods

### RNA-sequencing

Sample preparation was made using Illumina TrueSeq^TM^ RNA sample preparation kit with 1 µg of high quality total RNA and was sequenced using a paired end 101 bp protocol on the HiSeq2000 platform (Illumina). For INS1 cells, reads were aligned using Top Hat 2.0.0(Kim et al., 2013) and Bowtie version: 2.0.0.5 (Langmead and Salzberg, 2012) to the rat RGSC3.4 genome assembly. For human islet, 66 samples (all non-diabetic) were sequenced. Sequencing reads were aligned to the human reference genome (hg19) with STAR version 2.3.0e (Dobin et al., 2013) and counted using HTseq v0.5.4 and edgeR (Robinson et al., 2010). The raw sequence reads have been deposited at European Nucleotide Archive (ERA261116) and at GEO (GSE50398) for INS1 and human islets, respectively.

### Real-time quantitative PCR and Western blot

Total RNA was extracted using RNeasy Plus Mini Kit (Qiagen), cDNA was prepared using RevertAid^™^ First Strand cDNA Synthesis Kit (Thermo Scientific). QPCR was performed using TaqMan assays (Table S5). mRNA were measured 48 hours (INS1 cells) or 24 hours (human islets) after silencing. For human islets, dispersed human islet cells in 804G-conditioned medium coated dishes(Hammar et al., 2004) were used. Protein (15 – 80 µg) was separated by SDS-PAGE, transferred to nitrocellulose membranes and detection was made using SuperSignalWestFemto Chemiluminescent Substrate (Thermo Scientific). Protein expression was measured 72 hours or 7 days after silencing. Antibodies used can be found in Table S5.

### Statistical Analysis

Statistical analyses were performed in IBM^®^ statistics version 21 (IBM Corporation, Armonk, NY). Student’s t-test was used when two groups were compared and paired t-test was used for analysis of GSIS, GSPS and TIRFM data. p < 0.05 was regarded as significant and two-tailed p-values were calculated. False discovery rate (FDR) of 5% was used to correct for multiple comparisons. All results are given as mean ± standard error of the mean (SEM) unless otherwise stated.

### Data Access

ArrayExpress Accession: E-MTAB-2034

## Acknowledgements

This work has been supported through funding from the Swedish Research Council, the Knut and Alice Wallenberg foundation, the Wallenberg foundation, the Lundberg foundation, the Swedish Diabetes Research Foundation and the European Commission’s Seventh Framework Program fund, the UMAS fund, Novo Nordisk Foundation, Magus Bergvall’s Foundation, Craafoord Foundation, Tore Nilson’s Foundation, the Hjelt Foundation, EFSD/MSD, the United States Public Health Service and by the National Institutes of Health, and the Kovler Family Foundation. We give our most sincere thanks to Mar Gonzàles-Porta, Karl Bacos, Jens Lagerstedt, Anna-Maria Veljanovska Ramsay, Ulrika Krus, Jasmina Kravic, Lena Eliasson, Anders Rosengren, Emma Ahlqvist, Peter Osmark and Alex Persson (Lund University) for constructive suggestions and technical assistance. The human islet was provided from the Human Tissue Laboratory at Lund University in collaboration with Olle Korsgren (Uppsala University) and the Nordic Network for Clinical Islet Transplantation. The authors declare that there is no duality of interest associated with this manuscript.

